# Early detonation by sprouted mossy fibers enables aberrant dentate network activity

**DOI:** 10.1101/466300

**Authors:** William D. Hendricks, Gary L. Westbrook, Eric Schnell

## Abstract

In temporal lobe epilepsy, sprouting of hippocampal mossy fiber axons onto dentate granule cell dendrites creates a recurrent excitatory network. However, unlike mossy fibers projecting to CA3, sprouted mossy fiber synapses depress upon repetitive activation. Thus, despite their proximal location, large presynaptic terminals, and ability to excite target neurons, the impact of sprouted mossy fiber synapses on hippocampal hyperexcitability is unclear. We find that despite their short-term depression, single episodes of sprouted mossy fiber activation in hippocampal slices initiated bursts of recurrent polysynaptic excitation. Consistent with a contribution to network hyperexcitability, optogenetic activation of sprouted mossy fibers reliably triggered action potential firing in postsynaptic dentate granule cells after single light pulses. This pattern resulted in a shift in network recruitment dynamics to an “early detonation” mode and an increased probability of release compared to mossy fiber synapses in CA3. A lack of tonic adenosine-mediated inhibition contributed to the higher probability of glutamate release thus facilitating reverberant circuit activity.

**Significance Statement:** Sprouted mossy fibers are one of the hallmark histopathological findings in temporal lobe epilepsy. These fibers form recurrent excitatory synapses onto other dentate granule cells that display profound short-term depression. Here, however, we show that although these sprouted mossy fibers weaken substantially during repetitive activation, their initial high probability of glutamate release can activate reverberant network activity. Furthermore, we find that a lack of tonic adenosine inhibition enables this high probability of release and, consequently, recurrent network activity.

## Introduction

Mossy fibers contacting CA3 pyramidal cells have a low probability of release (P_r_), but show profound short-term facilitation (1). These “conditional detonator” synapses strongly drive post-synaptic cell firing during repetitive activation (2). In epilepsy, mossy fiber axons sprout collaterals onto the proximal dendrites of other dentate granule cells (3), where conditional detonation could be highly epileptogenic. However, sprouted mossy fibers rapidly depress during repetitive activation (4), which might limit the extent with which they could drive seizure activity (5).

Despite multiple lines of evidence demonstrating *de novo* recurrent connections in mouse models of epilepsy, the impact of mossy fiber sprouting on circuit dynamics remains uncertain (6–9). Sprouted mossy fibers are absent under non-pathological conditions, and the formation of novel recurrent connections capable of activating post-synaptic granule cells (10) could induce run-away excitation, particularly if they robustly activate the typically quiescent dentate network. Although increased dentate excitation occurs in temporal lobe epilepsy (11, 12), the challenge of isolating and selectively stimulating sprouted mossy fibers has severely hampered studies that directly examine sprouted mossy fiber synapses (10) and how the activation of these fibers might contribute to epileptiform activity.

Here, we selectively activated sprouted mossy fiber axons with optogenetics to determine their influence on post-synaptic granule cell firing and dentate gyrus activity. We find that sprouted mossy fibers reliably drive post-synaptic firing of dentate granule cells and initiate recurrent circuit activity. This effect is mediated by increased release probability at these synapses, attributed to a lack of tonic adenosine signaling in the inner molecular layer of the dentate gyrus. The increased P_r_, allowed sprouted mossy fiber activation to reliably recruit post-synaptic circuits, as primed network “spark plugs.”

## Results

As the functional contribution of mossy fiber sprouting to epileptogenesis remains controversial (5–8), we combined the pilocarpine model of temporal lobe epilepsy with electrophysiology to directly examine the contribution of sprouted mossy fiber activation to epileptiform activity (see methods). To isolate sprouted mossy fibers from other granule cell inputs, we used DcxCre::ChR2 mice to selectively label neonatally-born granule cells with channelrhodopsin2 (ChR2). We then prepared acute mouse hippocampal slices, and optogenetically activated sprouted mossy fibers while recording from ChR2-negative (unlabeled) granule cells (4, 13) (Fig. 1A-C; Supplementary Fig. 1). Optogenetic activation of sprouted mossy fibers evoked EPSCs in dentate granule cells only in slices from pilocarpine-treated mice (Fig. 1D,E; also see ref. 4). Postsynaptic cells, filled with Alexa568 dye during recordings, did not have hilar basal dendrites (0 of 32 cells; Fig. 1C), indicating that LED-evoked responses originated from recurrent (sprouted mossy fiber) synaptic inputs.

**Figure 1:**
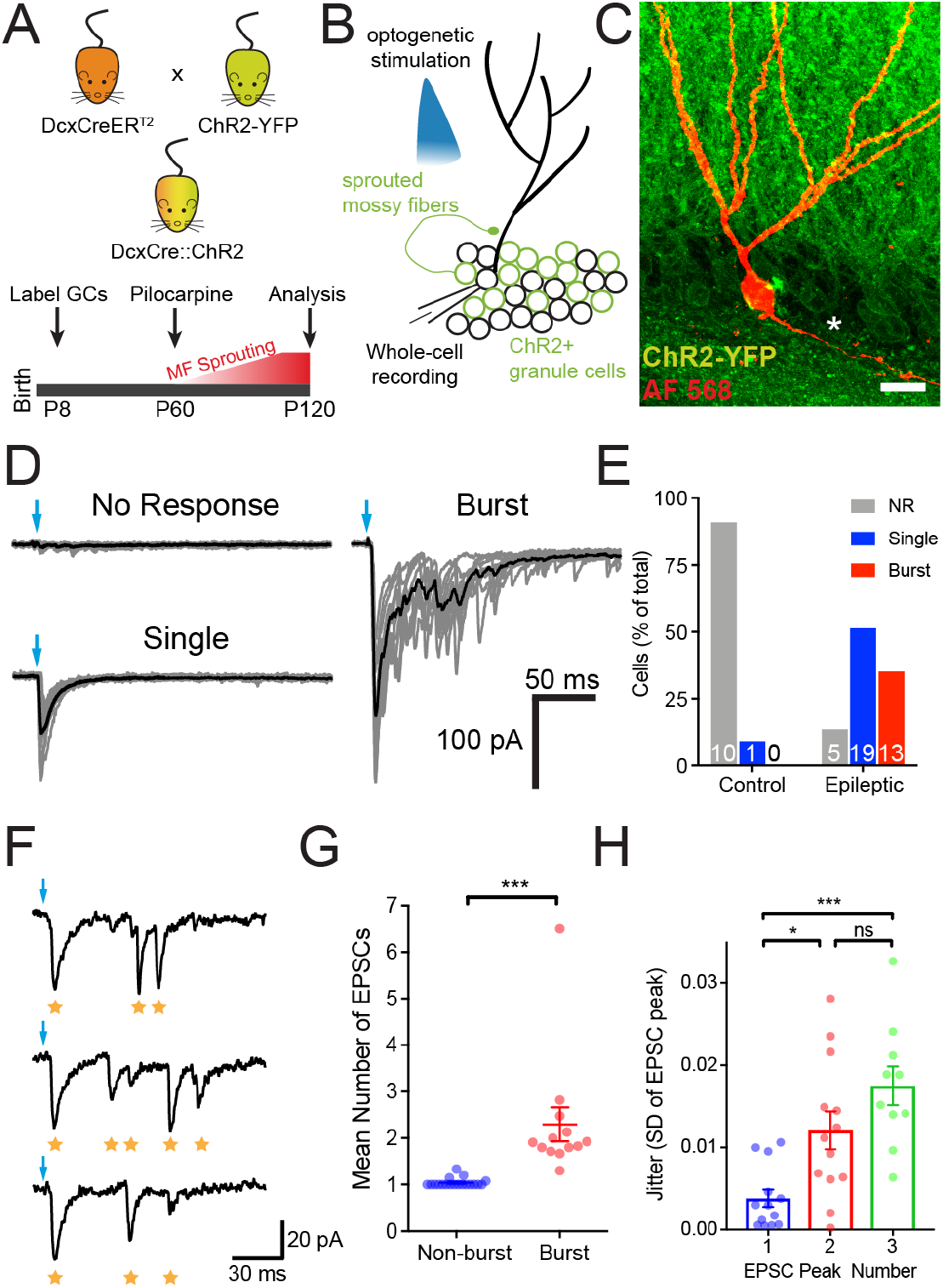
Stimulation of sprouted mossy fibers triggers spiking in post-synaptic dentate granule cells and recurrent network activity. (*A*) Experimental design for granule cell labeling and induction of epileptic mossy fiber sprouting. DcxCreER^T2^ mice were bred with conditional ChR2-YFP (Ai32) reporter mice (top) and given tamoxifen (TAM) at P8 to turn on reporter gene expression (bottom). At 2 months (P60), mice were given pilocarpine to induce seizures and mossy fiber sprouting or kept as controls. All analysis was at P120. (*B*) Whole-cell recordings were performed on ChR2-negative dentate granule cells and sprouted mossy fiber axons were stimulated with blue (470 nm) LED light delivered through the objective. (*C*) Representative confocal image of a recorded granule cell filled with AlexaFluor 568 (50 μM) during whole-cell recording (levels and gamma adjusted for clarity). Star designates mossy fiber axon; no basal dendrites were present (scale, 10 μm). (*D*) Example traces from cells with no response (top left), single EPSCs (bottom left), and epileptiform bursts (right) after single optogenetic stimulation of ChR2+ granule cells (blue arrows). Scale bar on right applies to all traces. (*E*) The frequency of cells with single EPSCs (Single), and EPSC bursts (Burst) in epileptic mice was greater than controls (Chi-square, p < 0.0001). Number of cells in each category are listed on the graph. (*F*) Three consecutive traces taken from a cell with EPSC bursts. Optogenetic stimulation (blue arrow) evoked an initial EPSC (likely monosynaptic) followed by variably timed burst EPSCs. Gold stars designate detected EPSCs. (*G*) Cells with bursts have greater average number of EPSC peaks (Mean number of peaks: Non-burst, n = 19; Burst, n = 13; unpaired t-test, t_30_ = 4.18, p = 0.0002). (F) Jitter (trial to trial variability of EPSC onsets, as standard deviation) taken from the first 3 peaks in an EPSC burst is increased after the first EPSC peak (EPSC jitter: Peak 1, n = 13; Peak 2, n = 13; Peak 3, n = 10; One-way ANOVA, F_2,33_ = 12.0, p = 0.0001; Tukey’s test, Peak 1 vs. Peak 2, p = 0.0105; Peak 1 vs. Peak 3, p < 0.0001; Peak 2 vs. Peak 3, p = 0.1544). High variability in second and third EPSC peaks within a burst suggest that they result from polysynaptic activity. Summary data presented as mean ± SEM. *p < 0.05, ***p < 0.001

In addition to monosynaptic sprouted mossy fiber–GC(smf-GC) EPSCs, we frequently observed LED-evoked epileptiform activity following single light pulses (Fig. 1D, right). These bursts were not a result of altered intrinsic properties compared to cells with single or no evoked recurrent EPSCs (R_i_, input resistance: cells without bursts, 610 ± 47 MΩ; cells with bursts, 527 ± 77 MΩ; unpaired t-test, t_30_ = 0.9711, p = 0.3392; C_m_, cell capacitance: non-bursts, 51.5 ± 3.8 pF; bursts, 46.8 ± 3.8; unpaired t-test, t_30_ = 0.8302, p = 0.4130). Optogenetic stimulation only elicited single spikes in ChR2-expressing granule cells (number of AP per 1 ms light pulse: 0.98 ± 0.09, n = 5 cells), indicating that sprouted mossy fiber activation initiated recurrent activity. Consistent with polysynaptic activation, EPSC peaks during bursts (EPSC peaks 2 – 3) had high jitter (Fig. 1 F-H). One possible source of these polysynaptic events is through the activation of additional populations of granule cells with their own sprouted mossy fibers, if sprouted mossy fibers would be able to effectively drive post-synaptic firing.

Unlike healthy mossy fiber – CA3 synapses (1) that are described as conditional detonators, sprouted mossy fiber synapses exhibit profound frequency-dependent short-term depression (Supplementary Fig. 2, see also ref. 4), which may limit their ability to drive postsynaptic firing (5). Thus, we compared patterns of synaptically-evoked spike generation in post-synaptic target cells at these two different synapses using a short 10 Hz train of optogenetic stimulation. In striking contrast to synaptic facilitation and delayed CA3 pyramidal cell spike generation at mf-CA3 synapses, 10 Hz stimulation of sprouted mossy fiber synapses induced post-synaptic spikes only at the beginning of the train (Fig. 2). Importantly, this firing pattern did not result from a failure to activate sprouted mossy fibers in epileptic brains, as these fibers maintain action potential fidelity throughout optogenetic trains (Supplemental Fig. 1). Thus, the failure to maintain post-synaptic activation resulted instead from reduced glutamate release later in the train, consistent with short-term synaptic depression at these synapses (4). Despite this depression, however, single pulses triggered circuit activation sufficient to recruit additional recurrent networks (Fig. 1D).

**Figure 2:**
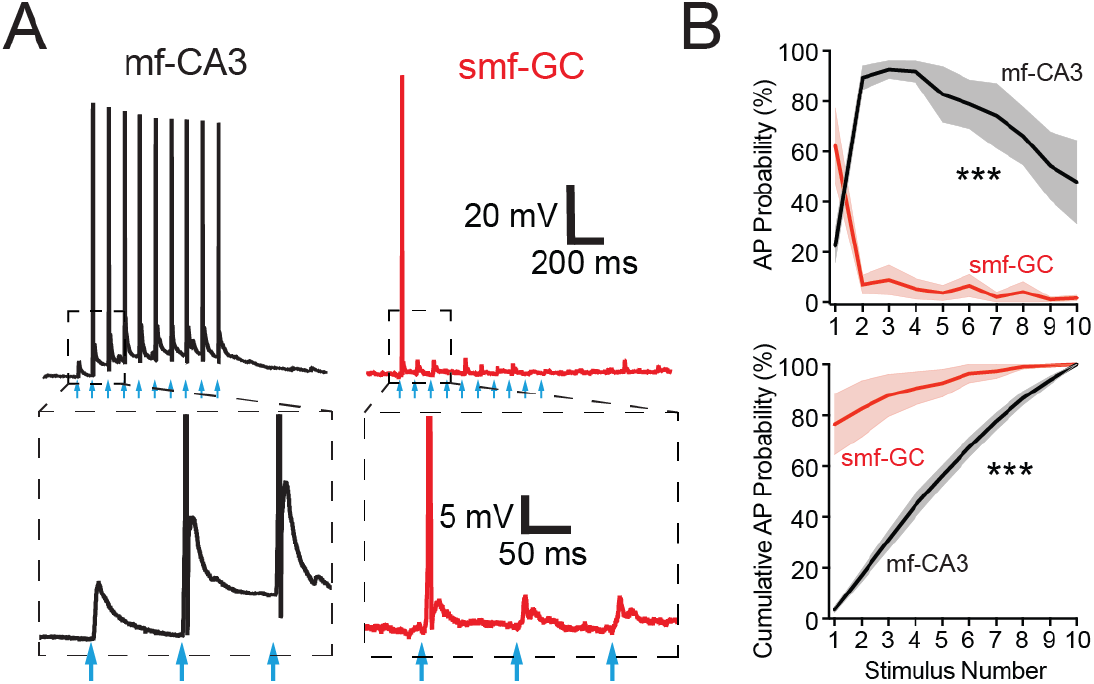
Sprouted mossy fiber activation drives post-synaptic granule cell firing early during brief trains. (*A*) Representative current clamp recordings during mossy fiber – CA3 (mf-CA3, black trace) and sprouted mossy fiber – granule cell (smf-GC, red trace) activation by 10 Hz, 10 pulse LED trains (blue arrows). Dashed boxes represent insets (below) highlighting facilitation of mf-CA3 EPSPs (left) and depression of smf-GC EPSPs (right). (*B*) Action potential (AP) probability (top) is shifted to the first stimulus after smf-GC activation relative to mf-CA3 activation (mf-CA3, n = 6 cells; smf-GC, n = 8 cells, 2-way RM ANOVA, F_1,12_ = 85.68, p < 0.0001; Sidak’s multiple comparisons test, Stimulus 1, p = 0.0048, Stimuli 2 – 9, all p < 0.0001, Stimulus 10, p = 0.0006). Number of cells in each category are listed on the graph. Summary data presented as mean ± SEM. ***p < 0.001

Both the frequency-dependent short-term depression and the robust initial recruitment of postsynaptic cell firing during recurrent activation (Fig. 2B) could result from a high initial probability of release (P_r_) at sprouted mossy fiber synapses. To directly test for increased P_r_, we measured use-dependent MK-801 block kinetics during activation of both types of mossy fiber synapses. Optogenetically-evoked NMDAR-mediated synaptic currents were isolated by voltage-clamping granule cells to -70 mV in Mg^2+^ free ACSF, in the presence of NBQX (10 μM) and SR95531 (10 μM) to block AMPA- and GABA_A_-receptors, respectively. After baseline EPSC recording, LED stimulation was paused for 10 minutes while MK-801 (40 μM) was washed onto the slice and allowed to equilibrate. Upon resuming LED stimulation (0.05 Hz), MK-801 progressively inhibited NMDAR-EPSCs from smf-GC and mf-CA3 synapses (Fig. 3A). The rate of decay was faster for sprouted mossy fiber synapses (Fig. 3B), indicating a higher P_r_. Interestingly, MK-801 block rate at smf-GC synapses was better fit by a double-exponential decay model (Fig. 3C), whereas at mf-CA3 synapses, a single exponential model was sufficient, suggesting increased synaptic heterogeneity at sprouted mossy fiber synapses (14). Overall, the increased P_r_ at sprouted mossy fiber synapses likely contributes to target granule cell firing and therefore the spread of recurrent excitation in epileptic brains.

**Figure 3:**
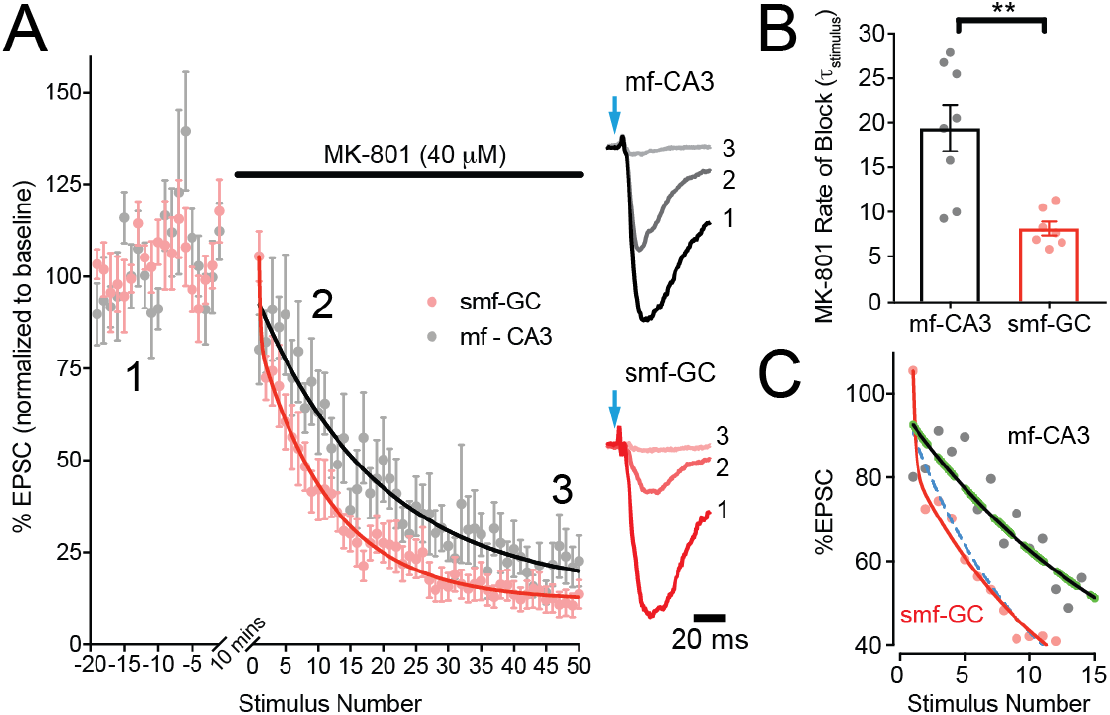
Elevated P_r_ at sprouted mossy fiber synapses. (*A*) Probability of release was measured from the kinetics of use-dependent MK-801 block of NMDAR-mediated EPSCs during 0.05 Hz LED stimulation. Averaged and normalized responses (left) are plotted before and after bath application of MK-801 (40 μM; mf-CA3, n = 8; smf-GC, n = 7). Representative, peak-scaled NMDAR EPSC averages (right) were taken during baseline, shortly after resuming LED stimulation, and at the end of the experiment, numbered 1, 2, 3, respectively. (*B*) Average rate of MK-801 block measured in individual cells. MK-801 rate of block was significantly faster at sprouted mossy fiber synapses (unpaired t-test, t_13_ = 3.942, p = 0.0017), indicating higher P_r_. (*C*) Same as in (*A*), but only showing the first 15 sweeps. Single (solid lines) and double (dashed lines) exponential fits are plotted for smf-GC (red and blue) and mf-CA3 (black and green). Sprouted mossy fibers are better fit by double exponential decays (extra sum of squares F-test, F_2,395_ = 5.07, p = 0.0067) whereas mf-CA3 synapses are fit identically by single and double models. All data are presented as mean ± s.e.m. **p < 0.01

Extracellular adenosine inhibits neurotransmitter release at healthy mossy fiber synapses in CA3 via A1-type adenosine receptors (A_1_Rs) located on mossy fiber boutons. This tonic activation of pre-synaptic A_1_Rs contributes to the profound short-term plasticity at these synapses (15, 16). As altered adenosine metabolism may be a contributing factor in epileptogenesis (17–19), we posited that sprouted mossy fibers might manifest an increased P_r_ due to a reduction in A_1_R-mediated inhibition. To measure tonic A_1_R-mediated inhibition of sprouted mossy fibers in epileptic mice, we washed on the selective high-affinity A_1_R antagonist, 8-Cyclopentyl-1,3-dipropylxanthine (DPCPX, 200 nM), while recording sprouted mossy fiber-mediated responses. In contrast to the enhancement of synaptic transmission at healthy mf-CA3 synapses (Fig. 4A,B) (15), DPCPX had no effect on sprouted mossy fiber EPSCs (Fig. 4A,B), indicating a lack of tonic A_1_R signaling. The DPCPX-induced increase in EPSC amplitudes at mf-CA3 synapses coincided with decreased PPR (paired-pulse ratio, P_2_/P_1_: pre-DPCPX, 2.36 ± 0.37; post-DPCPX, 1.37 ± 0.12, n = 6; paired t-test, p = 0.0191), as expected for a pre-synaptic effect on A_1_Rs.

**Figure 4:**
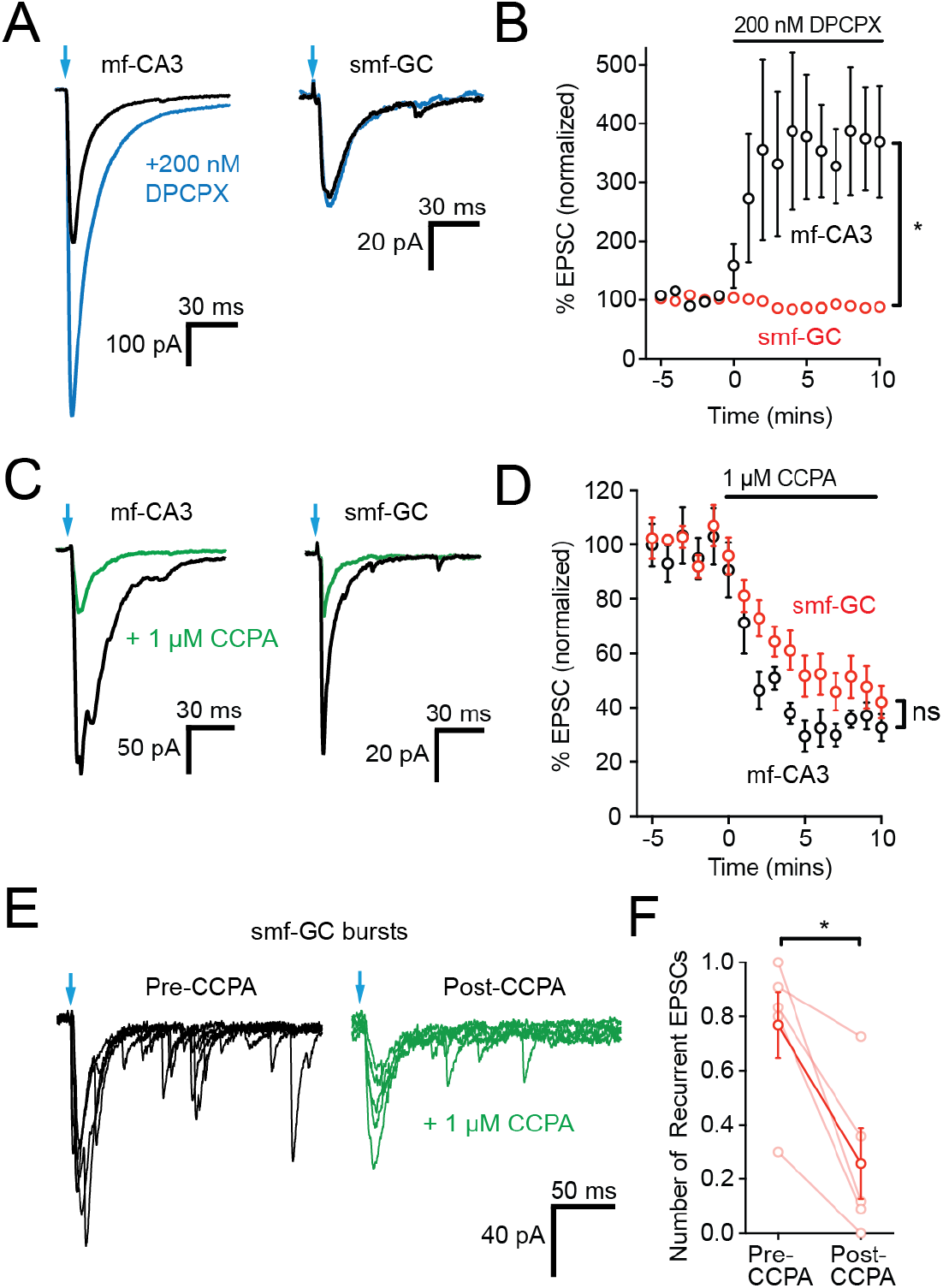
Lack of tonic adenosine contributes to increased release probability and polysynaptic activity. (*A*) Mossy fiber – CA3 (left) and sprouted mossy fiber-GC (right) EPSCs before (black traces) and after (blue traces) washing on the adenosine 1a receptor (A_1_R) antagonist DPCPX (200 nM). Bath application of DPCPX facilitates mf-CA3 EPSCs (n = 8 cells, paired t-test, *t*_7_ = 5.77, p = 0.0007), but has no effect on EPSC amplitudes at smf-GC synapses (n = 5 cells, paired t-test, *t*_4_ = 1.39, p = 0.2364). (*B*) Normalized EPSC amplitudes of mf-CA3 (black, n = 8 cells) and smf-GC (red, n = 5 cells) responses before and during DPCPX administration (unpaired t-test, *t*_11_ = 3.63 p = 0.0040). (*C*) The A_1_R agonist CCPA (1 μM) inhibits both mf-CA3 (left) and smf-GC (right) EPSCs similarly (baseline, black example traces; CCPA, green traces). (*D*) Normalized mf-CA3 (black) and smf-GC (red) EPSC amplitudes during CCPA application, demonstrating intact A_1_R responses at both synapses (mf-CA3, n = 7 cells, smf-GC, n = 12 cells; unpaired t-test, t_17_ = 1.08, p = 0.2954). (*E*) Adenosine A_1_R agonism inhibits recurrent polysynaptic network activity evoked by single pulse optogenetic stimulation of sprouted mossy fibers. Examples are 5 consecutive EPSC traces overlaid from the same cell, pre- and post-CCPA (black and green, respectively). (*F*) Quantification of EPSC burst inhibition by CCPA at smf-GC synapses. CCPA (1 μM) reduces the mean number of recurrent EPSCs during a burst (number of recurrent EPSCs per epoch, n = 5 cells, t_4_ = 3.963, p = 0.0166). Summary data presented as mean ± SEM. *p < 0.05

To determine whether the lack of tonic A_1_R-mediated inhibition at sprouted mossy fiber synapses resulted from an absence of presynaptic adenosine receptors, we washed on the selective A_1_R agonist, 2-Chloro-*N*^6^-cyclopentyladenosine (CCPA, 1 μM). CCPA reduced sprouted mossy fiber EPSCs (Fig. 4C,D) and increased paired-pulse facilitation (Supplementary Fig. 3). Thus A_1_Rs were present and functional on sprouted mossy fiber terminals, indicating that the lack of tonic A_1_R modulation was due to a reduced extracellular adenosine concentration. CCPA decreased the probability of release at sprouted mossy fiber synapses, but it did not restore frequency facilitation (Supplementary Fig. 4), suggesting that the molecular mechanisms underlying synaptic facilitation at mossy fiber terminals cannot be restored solely by increasing A_1_R activation.

Finally, to test whether reduced A_1_R activation and increased P_r_ at sprouted mossy fiber synapses contributes to recurrent circuit activation, we examined the effect of A_1_R activation using hippocampal slices from epileptic mice. When applied to slices demonstrating polysynaptic activity after sprouted mossy fiber activation, the A_1_R agonist CCPA reduced recurrent EPSCs as well as the charge transfer carried by polysynaptic bursts (Fig. 3E,F). Together, these data indicate that reduced A_1_R activation at granule cell outputs contributes to hyperexcitability in the dentate gyrus in epileptic brains.

## Discussion

Here we show that sprouted mossy fibers can trigger reverberating network activity despite their profound short-term depression. This recurrent network activation is enabled by a high probability of release, resulting in post-synaptic cell “early detonation”. Moreover, this is due, at least in part, to the lack of tonic adenosine inhibition at sprouted mossy fiber synapses in the inner molecular layer.

The retrograde sprouting of mossy fibers is common to both animal models of epilepsy and human patients with temporal lobe epilepsy, yet the functional impact of these recurrent synapses is not well understood (5, 6). We demonstrate that single optogenetic stimulation of these fibers can produce bursts of recurrent EPSCs, suggesting that sprouted mossy fibers effectively recruit the local network. This observation is consistent with paired granule cell recordings from epileptic mice (10), which suggest that sprouted mossy fibers form recurrent excitatory connections capable of driving post-synaptic cell spiking.

Notably, all triggered EPSC bursts were self-limited, which may be a manifestation of the strong short-term depression of this synapse (4, 10). This helps explain prior indirect observations made using extracellular retrograde stimulation of mossy fibers, which caused self-limited episodes of granule cell population spiking in slices from epileptic rats (20). We recognize that the conclusions that can be drawn using optogenetic stimulation in an acute slice preparation in regards to seizure activity *in vivo* are limited, primarily due to the isolation of slices from other brain networks and the cerebral extracellular environment. However, these recurrent networks were widespread within the 300 μm brain slices, demonstrated by our ability robustly trigger recurrent network activation, suggesting the extent of this recurrent network is likely an underestimate.

Nevertheless, we propose that increased P_r_ at sprouted mossy fibers provides a reasonable mechanism that could enable sprouted mossy fibers to act as “spark plugs” to hyperactivate local dentate gyrus networks. Our direct comparisons of relative P_r_ between smf-GC and mf-CA3 synapses are consistent with the larger success rate at individual smf-GC connections (0.35) (10) than the prior estimate of P_r_ at mf-CA3 synapses of 0.2-0.28 (21). We propose that the increased P_r_ at these sprouted synapses is, at least in part, due to reduced tonic inhibition by adenosine at the smf-GC synapse.

Does the lack of tonic adenosine in the dentate gyrus contribute to seizure activity in epilepsy? Although systemic or local injections of A_1_R agonists can reduce the severity of seizures (22, 23), adenosine has broad inhibitory effects throughout the brain. Here, the absence of tonic adenosine signaling at these synapses heightens the excitability of dentate gyrus circuitry, which was reduced by pharmacological A_1_R activation (Fig. 4). As the granule cell network is implicated in spontaneous seizures *in vivo* (24), restoration or augmentation of extracellular adenosine could reduce recurrent circuit dynamics in the in the dentate gyrus and provide one mechanism for the observed *in vivo* effects (22, 23). Thus, although seizure activity in epileptic brains likely results from multiple hyperexcitable circuit elements, the lack of tonic adenosine signaling in the dentate gyrus shifts sprouted mossy fiber synapses from “conditional detonators” to “early detonators”, which could contribute to the initiation of generalized seizures.

## Methods

### Animals

All experiments were carried out in accordance with local, state, and federal guidelines, and protocols were approved by the OHSU Institutional Animal Care and Use Committee. Housing was provided by Oregon Health & Science University’s Department of Comparative Medicine vivarium accredited by the Association for Assessment and Accreditation of Laboratory Animals. To generate DcxCre::ChR2 mice, homozygous *doublecortin-CreER^T2^* (line F18; RRID:MGI:5438982) mice were bred with homozygous *Gt(ROSA)26Sor^tm32(CAG-COP4*H134R/EYFP)Hze^* (Ai32; RRID:IMSR_JAX:012569) and used as previously described (4, 13). Briefly, Cre-mediated combination was induced by tamoxifen (TAM) at P8 (2 injections, 12 hours apart, 20 mg/kg in corn oil, i.p.), to permanently label neonatally-generated granule cells with ChR2-eYFP in DcxCre::ChR2 heterozygous mice. Status epilepticus was induced in two-month-old male mice with pilocarpine (325 mg/kg i.p.; Cayman Chemicals) after pre-treating with an i.p. injection of scopolamine methyl bromide (Sigma-Aldrich). Seizures were graded on the modified Racine Scale (25); status epilepticus (SE) was defined when a mouse had 3 or more Racine Grade 3 seizures, followed by continuous grade 2 seizing. Following 2 hours of SE, seizures were terminated with diazepam (10 mg/kg i.p.; Hospira, Inc.) and given soft food and i.p. injections of 5% glucose in 0.45% normal saline to aid in recovery. Mice that did not develop SE were humanely killed and excluded from further analysis.

### Slice Physiology

Acute brain slices for *ex vivo* electrophysiology were prepared as previously described (4). In brief, four-month-old male DcxCre::ChR2 mice were anesthetized with 4% isoflurane, followed by injection of 1.2% avertin (Sigma-Aldrich). Mice were transcardially perfused with 10 mL of ice-cold N-methyl-D-glucamine (NMDG)-based cutting solution, containing the following (in mM): 93 NMDG, 30 NaHCO3, 24 glucose, 20 HEPES, 5 Na-ascorbate, 5 N-acetyl cysteine, 3 Na-pyruvate, 2.5 KCl, 2 thiourea, 1.2 NaH2PO4, 10 MgSO4, and 0.5 CaCl2. Mice were rapidly decapitated, hippocampus dissected, and 300 μm transverse hippocampal sections were prepared in ice-cold NMDG solution with a Leica VT1200S vibratome. Hippocampal dissections were omitted for CA3 recordings and 300 μm sagittal slices were prepared to preserve pyramidal cells. Slices from both preparations recovered in warm NMDG cut solution for 15 mins followed by standard ACSF at room temperature for 1 hour prior to recording.

Dentate granule cell and CA3 pyramidal cell recordings were obtained with 3-5 MΩ borosilicate glass pipettes filled with a Cs^+^-based internal solution for voltage-clamp experiments, or a K^+^-based internal solution for current-clamp experiments. The Cs^+^-based internal solution contained the following (in mM): 113 Cs-gluconate, 17.5 CsCl, 10 HEPES, 10 EGTA, 8 NaCl, 2 Mg-ATP, 0.3 Na-GTP, 0.05 Alexa Fluor 568, pH adjusted to 7.3 with CsOH, with a final osmolarity of 295 mOsm; QX-314-Cl (5 mM; Tocris Bioscience) was included to block unclamped action potentials. The K^+^-based internal solution contained (in mM): 130 K-gluconate, 20 KCl, 10 HEPES, 4 Mg-ATP, 0.3 Na-GTP, 0.1 EGTA, 0.05 AlexaFluor 568, pH adjusted to 7.2 KOH, with a final osmolarity of 295 mOsm. Granule cells and CA3 pyramidal cells were identified with infrared differential interference contrast microscopy on an Olympus BX-51WI microscope. Whole-cell recordings were obtained by making high-resistance seals (>5 GΩ) and applying brief suction. Cells were filled with Alexa Fluor 568 dye to visually confirm cell type and assess for presence of hilar basal dendrites. Series resistance was uncompensated and cells with a >30% change in series resistance were excluded from analysis. Liquid junction potential was 8 mV and was uncorrected. For current-clamp recordings, minimal negative current was injected if necessary to maintain a resting potential of -70 mV in dentate granule cells and CA3 pyramidal cells.

Pulses of blue LED-powered (Thorlabs) light (1 ms, 470 nm, 8 mW/cm^2^, 0.05 Hz) were delivered through a 40x water immersion objective, targeted at the stratum lucidum for CA3 recordings and inner molecular layer for granule cell recordings. Stimulation frequency was modified for various experiments as noted in the text. Signals were amplified with an AxoPatch 200B amplifier (Molecular Devices), low-pass Bessel-filtered at 5 kHz, and digitized and sampled at 10 kHz using a NIDAQ (National Instruments) analog-to-digital board. Data were captured using a custom Igor Pro 8 (Wavemetrics) script and NIDAQmx (National Instruments) plugins. For presentation of EPSCs, a 2 kHz Gaussian filter was applied, *post-hoc*.

### Statistical Analysis

Curve fitting and EPSC trace averaging was carried out in Igor Pro 8 (Wavemetrics) using built-in and custom functions, respectively. Epileptiform burst activity (multiple EPSCs to single stimulus) was determined by eye, aided by fitting a single exponential curve to the initial decay and identifying delayed EPSC peaks rising above the decay fit line. Peak detection was implemented in Igor Pro by identifying zero-point crossings of thresholded peaks on the first-order derivative of the EPSC. Additional statistical analysis was performed in Prism 8 (GraphPad). Normality was tested with the Shapiro-Wilk normality test prior to statistical test selection. Paired and unpaired t-tests were used for normally distributed datasets; Mann-Whitney and Wilcoxon matched-pairs signed rank test were used for non-parametric datasets. For all experiments, significance was determined by p < 0.05 (*p <0.05, **p <0.01, ***p < 0.001). All summary data is presented as mean ± SEM.

## Acknowledgments

We wish to thank Drs. Zhi-Qi Xiong and Xuewen Cheng (Shanghai Institute for Neuroscience) for providing the DcxCreER^T2^ mouse line, and members of the Schnell and Westbrook laboratories for critical feedback and discussion on the manuscript. Research funding was provided by Department of Veterans Affairs, Veterans Health Administration, Office of Research and Development, Biomedical Laboratory and Development CDA-2 award 005-10S (ES); Department of Veterans Affairs Merit Review Award I01-BX002949 (ES); a Department of Defense CDMRP Award W81XWH-18-1-0598 (ES); a National Institutes of Health (NIH) Grant F31-NS098597 (WDH); NIH Grant R01-NS080979 (GLW); the Ellison Medical Foundation (GLW); and NIH Grant P30-NS061800 (Oregon Health and Science University Advanced Light Microscopy Core). The contents of this manuscript do not represent the views of the U.S. Department of Veterans Affairs or the United States government.

## Supplementary Figures

**Supplementary Figure 1:**
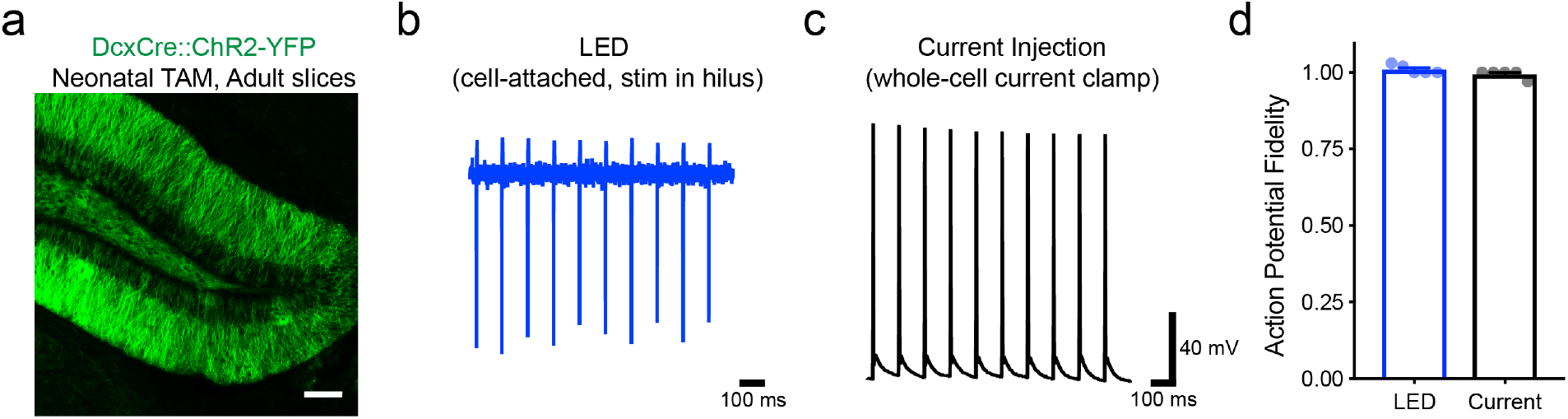
Action potential fidelity at 10 Hz stimulation in epileptic mice. a) Single confocal image from DcxCre::ChR2 mouse given tamoxifen neonatally (P6) and perfused at 4 months, demonstrating mature granule cell labeling. Scale bar, 100 μm. b) Example trace from a cell-attached recording of a ChR2 expressing dentate granule cell from an epileptic mouse reliably firing action potentials in response to optogenetic stimulation at 10 Hz (1 ms LED pulse, 470 nm, 8 mW/cm^3^). LED light was delivered via a 40x objective placed over the hilus to trigger antidromic action potential propagation and avoid direct stimulation of ChR2-expressing cell bodies. c) Example trace from a whole-cell current clamp recording of a dentate granule cell from an epileptic mouse, demonstrating that these cells reliably fire actions potentials during repeated brief current injections (1 ms, 500 pA, 10 Hz). d) Summary of action potential fidelity upon optogenetic (blue; n = 5) or current-injection (black; n = 5) stimulus trains. Actual potential fidelity potted as number action potentials evoked per LED stimulus. Data plotted as mean ± SEM.

**Supplementary Figure 2:**
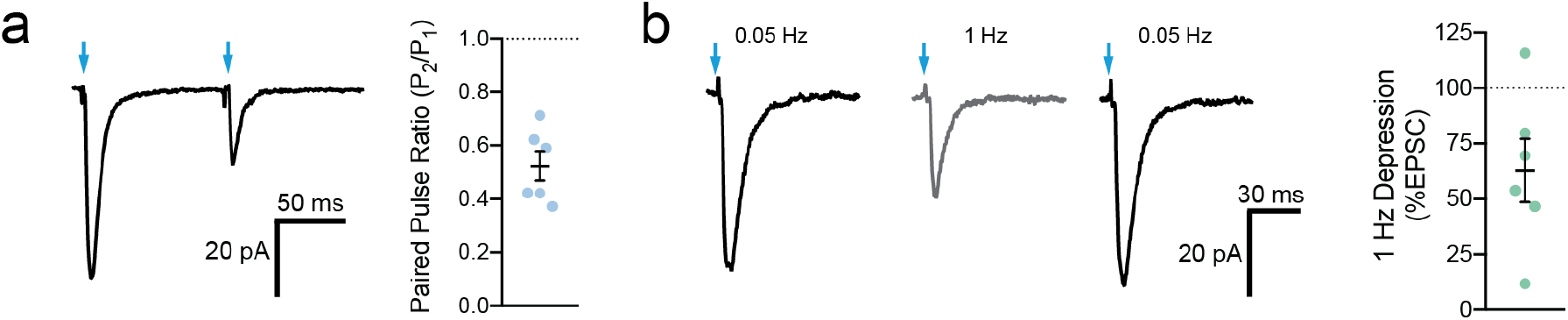
Sprouted mossy fibers exhibit paired-pulse depression and 1 Hz stimulation. (a) Example trace of paired pulse stimulation (100 ms ISI) of sprouted mossy fibers axons (left) and summary plot (right). Sprouted mossy fibers have a low paired-pulse ratio (PPR: 0.523 ± 0.06, n = 6). Blue arrows mark optogenetic stimulation. (b) Example average EPSC responses during baseline (0.05 Hz, left trace), 1 Hz stimulation (middle trace), and recovery (0.05 Hz, right trace). Summary data is plotted on the right. Sprouted mossy fibers display prominent depression during 1 Hz stimulation (%EPSC of baseline: 62.9 ± 14.3%, n = 6). Blue arrows mark optogenetic stimulation.

**Supplementary Figure 3:**
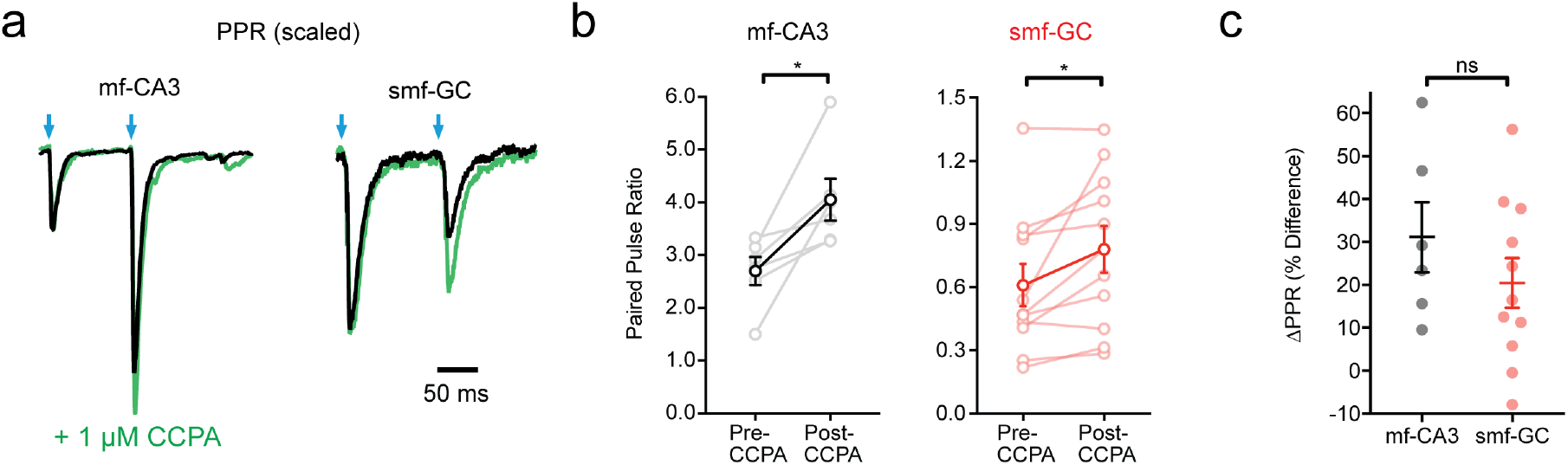
The A_1_R agonist CCPA increases the paired pulse ratio at both mossy fiber – CA3 and sprouted mossy fiber – GC synapses. (a) Representative EPSCs from paired-pulse experiments (100 ms ISI) from both mf-CA3 (left) and smf-GC (right) recordings before (black traces) and after (green traces) bath application of CCPA (1 μM, A_1_R agonist). Traces are scaled and normalized to the first EPSC peak. Larger relative EPSC peaks on the second pulse in CCPA indicates an increased paired-pulse ratio (PPR). (b) Quantification of paired-pulse experiments exemplified in (a). CCPA increases the PPR in both mf-CA3 (left) and smf-GC (right) experiments (PPR, mf-CA3: n = 6, paired t-test, t_5_ = 3.195, p = 0.0241; PPR, smf-GC: n = 11, paired t-test, t_10_ = 2.74, p = 0.0208). Group averages are overlaid over individual experiments. (c) Change in PPR from bath application of CCPA (% Difference) is similar for mf-CA3 and smf-GC synapses (ΔPPR: mf-CA3, n = 6; smf-GC, n = 11, unpaired t-test, t_15_ = 1.082, p = 0.2962).

**Supplementary Figure 4:**
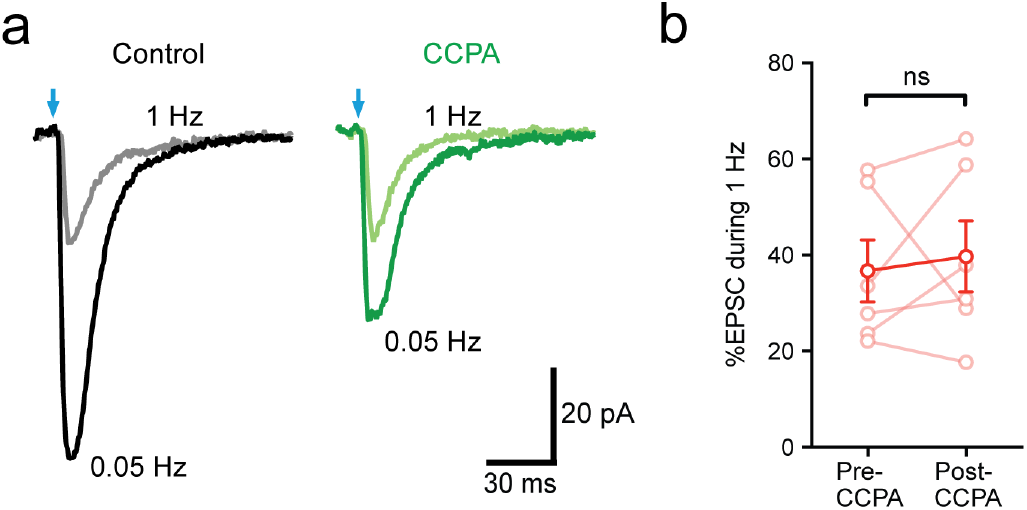
Restoring A_1_R tone with CCPA does not restore frequency-dependent facilitation at sprouted mossy fiber synapses. (a) Example traces taken at baseline stimulation frequency (0.05 Hz, darker traces) and during 1 Hz frequency trains (lighter traces) before (black traces) and after (green traces) bath application of CCPA (1 μM), demonstrating decreased EPSC amplitudes when evoked at 1 Hz. Examples are taken from the same cell/experiment. Scale bar is the same for both examples. (b) Quantification of experiments shown in (a). Bath application of CCPA does not alter EPSC depression during 1 Hz LED trains (% EPSC during 1 Hz, 50 pulse trains: pre-CCPA, n = 6; t_5_ = 0.422, paired t-test, p = 0.6905).

